# BpOmpW Antigen Stimulates the Necessary Immune Correlates of Protection Against Melioidosis

**DOI:** 10.1101/2021.05.16.444297

**Authors:** Julen Tomás-Cortázar, Lorenzo Bossi, Conor Quinn, Catherine Reynolds, David Butler, Niamh Corcoran, Maitiú Ó Murchú, Eve McMahon, Mahavir Singh, Patpong Rongkard, Juan Anguita, Alfonso Blanco, Susanna J. Dunachie, Danny Altmann, Rosemary Boyton, Johan Arnold, Severine Giltaire, Siobhán McClean

## Abstract

**SUMMMARY:** Melioidosis is a fatal disease caused by *Burkholderia pseudomallei* Gram-negative bacteria. It is the causative of 89,000 deaths per year in endemic areas of Southeast Asia and Northern Australia. Diabetes mellitus is the most risk factor, increasing 12-fold the susceptibility for severe disease. IFN-γ responses from CD4 and CD8 T cells, but also from NK and NKT cells are necessary to eliminate the pathogen. Elucidating the immune correlates of protection of our previously described protective BpOmpW vaccine is an essential step of any vaccine before clinical trials. Thus, we immunized non-insulin resistant C57BL/6j mice and an insulin resistant C57BL/6j mouse model of Type 2 Diabetes (T2D) with BpOmpW using Sigma Adjuvant System (SAS) (treatment) or SAS only (control). Two weeks later bloods and spleens were collected and serological analysis & in vitro exposure of splenocytes to the antigen for 60 hours were performed in both controls and treatment groups to finally analyze the stained splenocytes by flow cytometry. BpOmpW induced strong antibody response, stimulated effector CD4^+^ and CD8^+^ T cells and CD4^+^ CD25^+^ Foxp3^+^regulatory T cells and produced higher IFN-γ responses in CD4^+^, CD8^+^, NK and NKT cells relative to the control group in non-insulin resistant mice. T cell responses of insulin resistant mice to BpOmpW were comparable to those in non-insulin resistant mice. In addition, as a precursor to its evaluation in human studies, humanised HLA-DR and HLA-DQ transgenic mice elicited IFN-γ recall responses in an ELISPoT-based study and PBMCs from donors that were in contact to BpOmpW for seven days experienced T cell proliferation. Finally, plasma from melioidosis survivors with diabetes recognized our BpOmpW vaccine antigen. Overall, these range of approaches used strongly indicate that BpOmpW elicits the required immune correlates of protection to combat melioidosis and bring the vaccine closer to clinical trials.

## INTRODUCTION

Melioidosis is a potentially fatal tropical infection caused by the Gram-negative facultative intracellular bacillus *Burkholderia pseudomallei.* Melioidosis is endemic but increasingly emerging throughout the tropics. The global incidence is estimated to be 165,000 cases per year with 89,000 deaths and a significant global burden in terms of death and quality of life (Limmathurotsakul et al 2016; Birnie et al, 2019). Infections generally arise from environmental exposure and present as a spectrum of disease ranging from local pathologies such as pneumonia or abscesses to systemic disease and sepsis (White, 2003; Wiersinga et al., 2008; Wiersinga et al., 2018). The case fatality rate varies from 35 to 42 % in Thailand (Hinjoy et al., 2018) to 26% recorded in Australia (Stephens et al., 2016). Importantly, individuals with diabetes mellitus (DM) have a 12-fold increased susceptibility to melioidosis and experience more severe disease (Wiersinga et al., 2018). DM affects over 450 million people worldwide (Cho et al. 2018) of which, 90% are considered to have Type 2 Diabetes DM (T2DM) (Saeedi et al., 2019) and more than 50% of these individuals live in melioidosis endemic regions in Southeast Asia and Northern Australia (Dunachie & Chamnan, 2019). There is no licensed vaccine available to protect people in endemic regions from melioidosis, including those with T2DM.

We previously reported the identification of an efficacious antigen against melioidosis, identified using a proteomic approach based on the homology between *B. pseudomallei* and *B. cenocepacia* complex (Bcc) (Casey et al., 2016; McClean et al., 2016). We showed that the protective Bcc homologue OmpW in *B. pseudomallei* (BpOmpW) protected two different mouse models, BALB/c and C57BL/6J, from *Bp* challenge with distinct MHC haplotypes against melioidosis (Casey et al. 2016). In particular, we showed that 75% of immunised mice survived a lethal infection for an extended period of 81 days, a sustained protection not previously shown for any single subunit vaccine and surpassing that of the live attenuated vaccine 2D2, the benchmark against which all melioidosis vaccines are compared.

Understanding the correlates of protection is an important step in the development of any vaccine. Although the correlates of protection against melioidosis are poorly understood, it is clear that protection requires competent cellular immune responses mediated by T lymphocytes in both mice (Ketheesan et al. 2002) and humans (Jenjaroen et al., 2015). In particular, elevated IFN-γ responses associated with CD4 and CD8 T cells are essential to combat the disease (Ketheesan et al., 2002). However, IFN-γ-producing NK and NKT cells also participate in the response against melioidosis in mice (Haque et al., 2006) and in humans (Kronsteiner et al., 2019; Rongkard et al., 2020). Moreover, humoral immunity also contributes to the elimination of the bacteria in both mice and humans (Healey et al, 2005; Chaichana et al., 2018).

In order to further evaluate the BpOmpW antigen with a view to human trials, we have undertaken an investigation to establish the immune correlates of protection for this vaccine antigen. In particular, we have performed an in-depth analysis of the T cell responses associated with the BpOmpW antigen in C57BL/6J mice. Moreover, as diabetes is the most important risk factor for severe disease, most likely due to immune function dysregulation (Graves & Kayal et al. 2008; Daryabor et al., 2019), we also developed an insulin resistant mouse model to evaluate the immune responses to the BpOmpW antigen in the context of diabetes as recommended by the Steering Group on Melioidosis Vaccine development (Limmathurotsakul et al., 2015). Finally, we have examined the responses of human peripheral blood mononuclear cells (PBMCs) to the antigen.

## RESULTS

### BpOmpW activated T cells and induced effector CD4^+^ and CD8^+^ T cells

Activation of T cells is particularly important for effective vaccines against intracellular pathogens (Gilbert, 2012). Furthermore, given that T cell responses, especially IFN-γ responses, contribute to survival in acute melioidosis patients (Jenjaroen et al., 2015), we examined the T cell responses associated with BpOmpW immunization. Groups of 11 mice were immunized once with recombinant BpOmpW. Serological analysis of the mouse sera following immunization showed strong seroconversion (Total IgG, IgG1 and IgG2a) at two weeks despite the mice receiving only one immunization (Fig. 1, A; Fig. S1, A-C). T cell responses were then determined in splenocytes that had been re-exposed (treatment) to antigen *in vitro* by measuring cytokine responses and a range of T cell markers using flow cytometry compared to antigen exposed splenocytes from the adjuvant only group (SAS control), (Fig. 1, A). Activation of BpOmpW re-stimulated splenocytes was demonstrated by a significant decrease in CD45RB expression (p < 0.0001; Fig. 1, B), while the levels of CD25 and CD44 were significantly increased in response to BpOmpW re-exposure in both CD4^+^ and CD8^+^ T cells when compared to splenocytes from SAS control mice (p < 0.0001; Fig. 1, C-F). To further analyse the activation of T cells in immunized mice, we evaluated the relative expression of CD45RB and CD44 in both CD4^+^ and CD8^+^ T cells. BpOmpW stimulated the differentiation of T cells from naïve to effector T cells in the immunized mice compared to controls. Effector CD4^+^ CD45RB^lo^ CD44^hi^ T cells were over-represented in antigen re-exposed splenocytes (p < 0.0001; Fig. 1, G). Consistent with this, the naïve CD4^+^ CD45RB^hi^ CD44^lo^ subset was significantly reduced in the BpOmpW immunized group relative to SAS control splenocytes (p < 0.0001; Fig. 1, H). Likewise, both effector CD8^+^ T cells CD45RB^hi^ CD44^hi^ (Fig. 1, I) and CD45RB^lo^ CD44^hi^ (Fig. 1, J) were increased following BpOmpW immunization compared to SAS controls, whereas the number of naïve CD45RB^hi^ CD44^lo^ CD8^+^ T cells were reduced (Fig. 1, K). These data show that the immunization with BpOmpW results in the generation of strong CD4^+^, CD8^+^ T and B cell responses.

**Figure 1.**
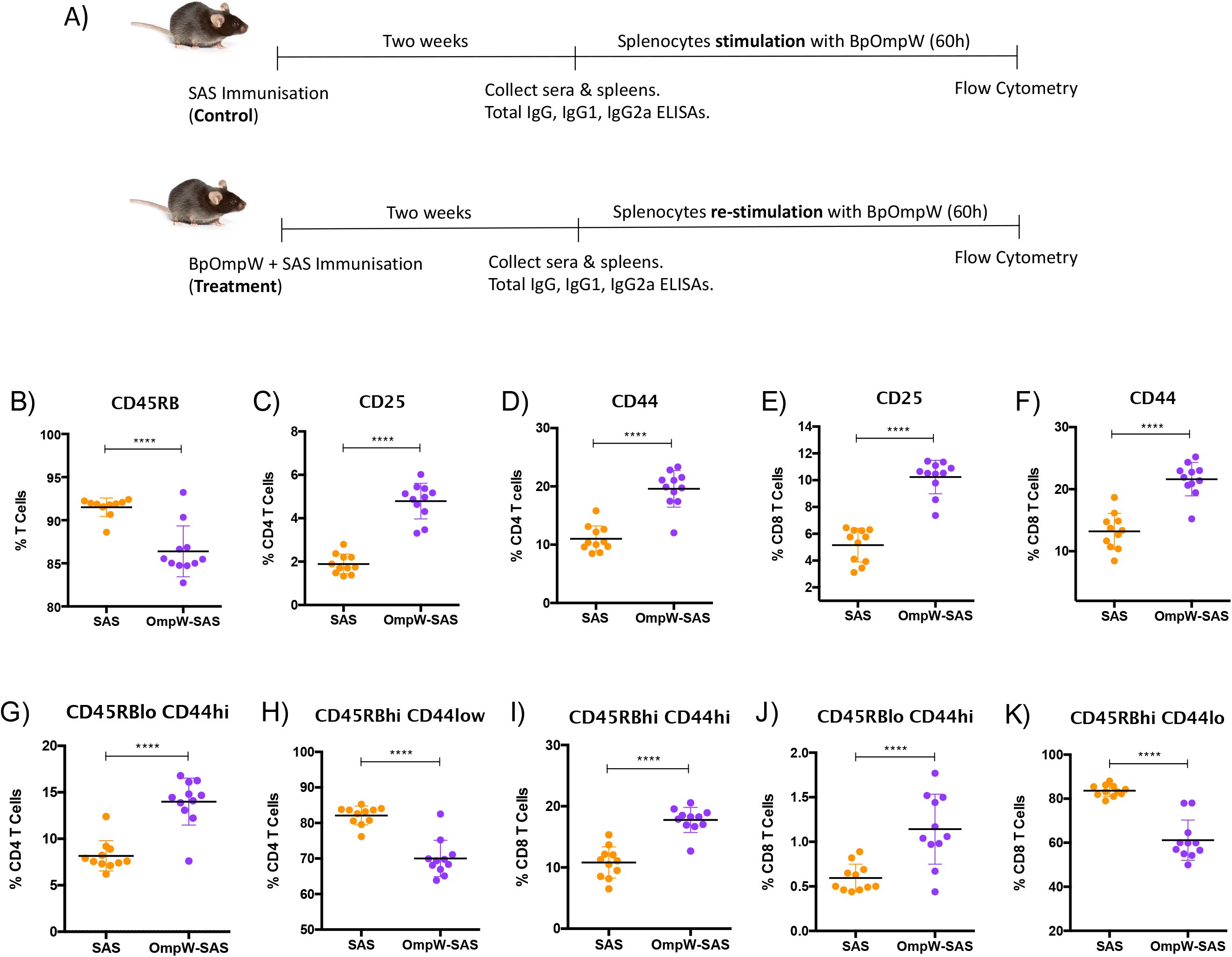
BpOmpW activated T cells and induced effector CD4^+^ and CD8^+^ T cells. A) Schematic illustration of the experimental timeline of non-insulin resistant mouse study. B) Percentages of parent cells expressing CD45RB. C to F) Percentages of CD4 and CD8 T cells expressing CD25 and CD44 activation markers. G to K) Percentages of different populations of CD4 and CD8 T cells defined by different levels of CD45RB and CD44 such as (G) effector CD4 T cells CD45RBlow CD44high, (H) naïve CD4 T cells CD45RBhigh CD44low, (I) effector CD45RBhigh CD44high CD8 T cells, (M) effector CD8 T cells CD45RBlow CD44high and (K) naïve CD8 T cells CD45RBhigh CD44low. SAS: splenocytes from adjuvant only immunised mice exposed to BpOmpW (Orange circles, Control). OmpW-SAS: splenocytes from SAS adjuvanted BpOmpW immunized mice re-exposed with BpOmpW in vitro (Purple circles, Treatment). Asterisks denote statistically significant differences according to two-tailed t-test. The significant levels are represented as follows: (p < 0.05, *); (p < 0.01, **); (p < 0.001, ***) (p < 0.0001, ****).

We also investigated the CD8+ and CD4+ T cell populations more in detail and we observed that the BpOmpW immunisation induced the appearance of CD8^hi^ and CD4^hi^ populations (Fig. 2, A) that were absent in the control group splenocytes (Fig. 2, B). Further analysis of these populations demonstrated that they were predominantly CD8^hi^ CD45RB^hi^ CD44^lo^ (Fig. 2, C; in red) and CD4^hi^ CD45RB^hi^ CD44^lo^ subpopulations (Fig. 2, D; in blue), in contrast to SAS control splenocytes that did not show these subpopulations (Fig. 2, E-F). Furthermore, splenocytes from SAS control mice showed a CD4^hi^ population (Fig. 2, B), to a lesser extent than BpOmpW group, although the population was predominantly CD45RB^lo^ CD44^hi^ (Fig. 2, F).

**Figure 2.**
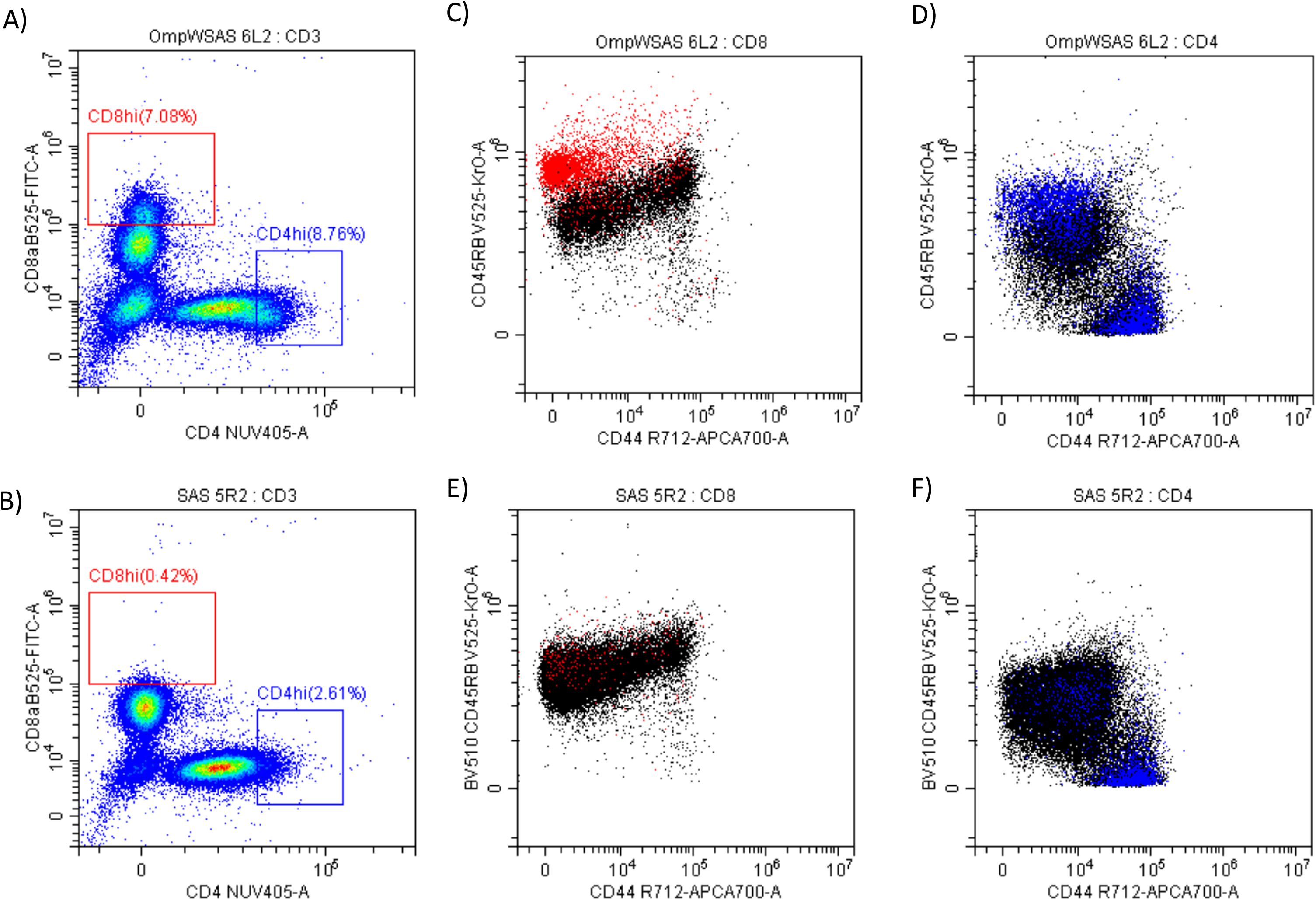
BpOmpW stimulated CD8hi and CD4hi populations that are CD45RBhi CD44lo. A-B) CD8hi (in red) and CD4hi (in blue) populations splenocytes from both SAS adjuvanted BpOmpW immunized mice re-exposed to BpOmpW (A) and from adjuvant only immunised mice exposed to BpOmpW (B). C-D) CD4 (C) and CD8 (D) T cell co-expression of CD45RB and CD44 in BpOmpW re-exposed splenocytes from SAS adjuvanted BpOmpW immunized mice. E-F) CD4 (E) and CD8 (F) T cell co-expression of CD45RB and CD44 in BpOmpW exposed splenocytes from SAS adjuvanted saline immunized mice.

### BpOmpW immunisation stimulated IFN-responses in CD4^+^, CD8^+^, NK, and NKT cells and increased regulatory T cells

IFN-γ responses dominate the immune response induced by melioidosis infection in patients (Koh et al., 2013). Therefore, in order to elucidate the T cell responses associated with BpOmpW immunisation, we evaluated a range of cytokine levels by flow cytometry. Although the autocrine growth factor IL-2 did not change in CD4^+^ T cells, it was upregulated in CD8^+^ T cells from the BpOmpW immunized group relative to the SAS control group (Fig. 3, A-B). The antigen elicited high levels of IFN-γ, IL-4 and IL-17, indicating that CD4^+^ T cells differentiated to Th1, Th2, and Th17 cells, indicative of a mixed Th response being elicited by BpOmpW (p < 0.0001, p < 0.0001, p < 0.0001, respectively; Fig. 3, C-E). Additionally, IFN-γ producing CD8^+^ T cells were also more abundant following BpOmpW immunisation (p = 0.011, Fig. 3, F). NK and NKT cells have been associated with patient responses to melioidosis, consequently we evaluated the responses of NK and NKT cells in BpOmpW-re-exposed splenocytes. A population of T cells that were negative for both CD4 and CD8 i.e. double negative (DN) cells were elevated in the BpOmpW immunized group relative to the SAS control group (p = 0.0001; Fig. 3, G). Expression of CD49b+ indicated that NKT cells constituted virtually all these DN cells (Fig. 3, H). Moreover, IFN-γ and IL-17 responses produced by NKT cells were increased in BpOmpW immunized mice in comparison with the SAS adjuvanted control mice in contrast to IL-4 which remained comparable to SAS control splenocytes (p = 0.0062, p < 0.0001, p = 0.942, respectively; Fig. 3, I-K). Finally, IFN-γ-producing CD49b^+^ NK cells were also significantly higher in splenocytes from BpOmpW immunized mice following re-exposure to antigen, relative to the adjuvant only control group (p = 0.0025; Fig. 3, L).

**Figure 3.**
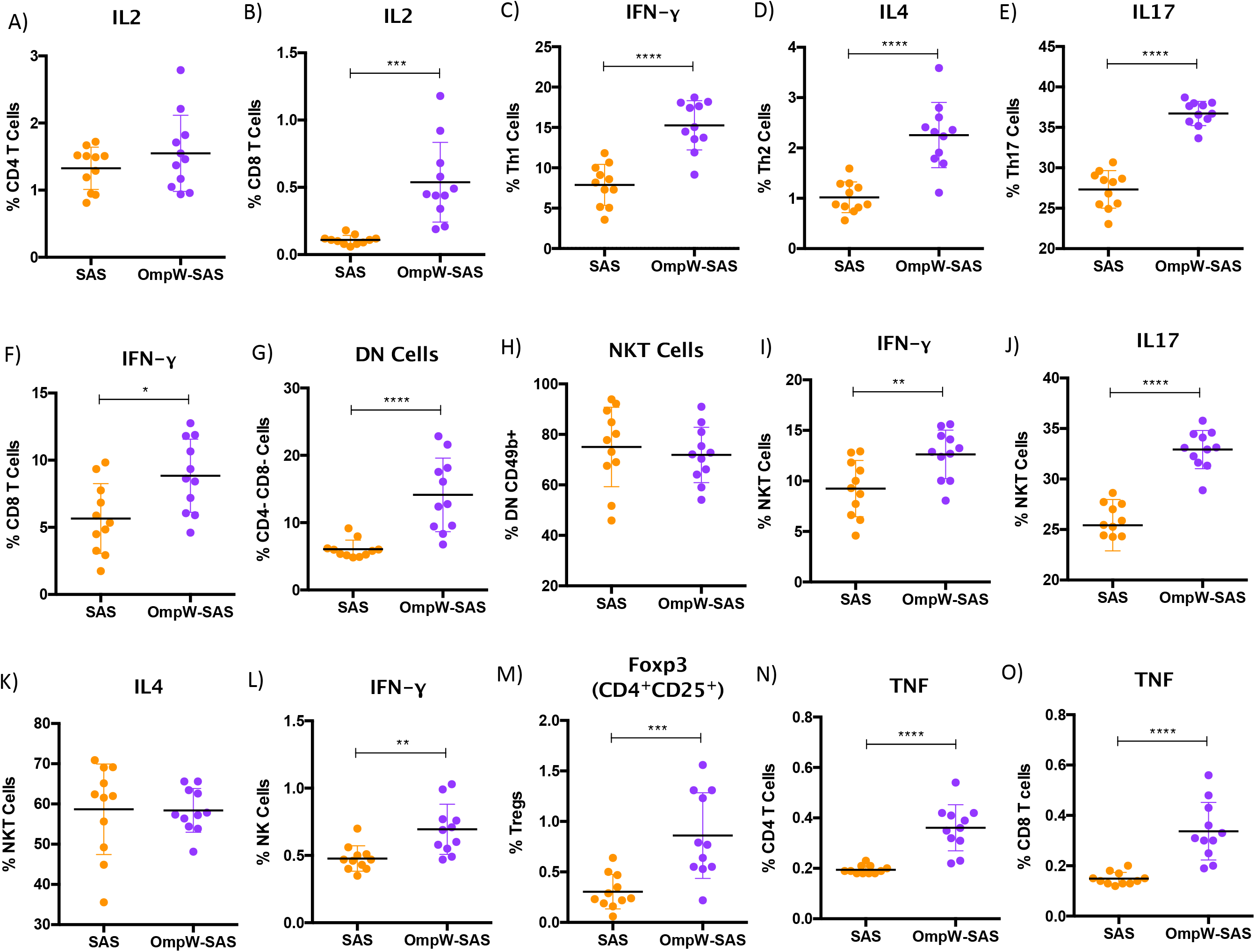
BpOmpW immunisation stimulated IFN-γ responses in CD4^+^, CD8^+^, NK and NKT cells and upregulated regulatory T cells. A-B) Percentages of CD4 (A) and CD8 (B) T cells expressing IL-2 cytokine. (C-E) Percentages of CD4 T cells expressing IFN-y (C), IL-4 (D) and IL-17 (E). G) Percentage of Double Negative (DN) cells (CD4^−^ CD8^−^). H) Percentage of Natural Killer T cells (NKT). (I-K) Percentages of NKT cells expressing IFN-y (I), IL-17 (J) and IL-4 (K). L) Percentages of Natural Killer (NK) cells expressing IFN-y. M) Percentages of Regulatory T cells (Tregs). N-O) Percentages of CD4 (N) and CD8 (O) T cells expressing TNF. SAS: splenocytes from adjuvant only immunised mice exposed to BpOmpW (Orange circles, Control). OmpW-SAS: splenocytes from SAS adjuvanted BpOmpW immunized mice re-exposed with BpOmpW in vitro (Purple circles, Treatment). Asterisks denote statistically significant differences according to two-tailed t-test. The significant levels are represented as follows: (p < 0.05, *); (p < 0.01, **); (p < 0.001, ***) (p < 0.0001, ****).

We also examined regulatory T cells as they can suppress proinflammatory damage produced by bacterial infections. Regulatory T cells were augmented in response to BpOmpW re-exposure in immunized mice relative to the spleen cells from adjuvant only control mice (p = 0.0007; Fig. 3, M). Finally, many studies have observed elevated expression of TNF during melioidosis infection in humans and animal models (Wiersinga et al., 2007; Krishnananthasivam et al., 2017). In this study, TNF was upregulated in both CD4^+^ and CD8^+^ T cells from BpOmpW immunized mice (p < 0.0001, p < 0.0001, respectively; Fig. 3, N-O).

### The immune response of insulin resistant mice to BpOmpW was predominantly comparable to that of non-insulin resistant mice

Due to the exquisitely enhanced susceptibility of people with diabetes to melioidosis infection, and that type 2 diabetes (T2D) is on the rise in tropical and subtropical regions, we needed to understand the immune response in the context of diabetes. We developed a polygenic insulin resistant mouse model by feeding C57BL/6J male mice with a high-fat diet (HFD) for up to 16 weeks as a model of T2D diabetes. Mice on the HFD continuously gained more weight than their litter mates on a normal diet from the first two weeks of HFD feeding (25.49 ± 2.12 vs 29.38 ± 0.82; p = 0.00001) at week 12 (Fig S2 B). Moreover, starting at week eight, HFD-fed mice began to develop hyperglycemia (HG: 13.17 ± 2.16 vs 15.66 ± 2.77; p = 0.02) and insulin resistance (IR (t_45_): 46.52 ± 16.26 vs 59.89 ± 23.20; p = 0.13), the latter more apparent at 12 weeks of treatment with the diet (HG: 11.44 ± 1.89 vs 13.82 ± 3.11; p = 0.044. IR (t_45_): 44.19 ± 8.20 vs 98.32 ± 17.92; p < 0.00001) (Fig S2 C-E). In addition, large lipid droplets were observed in the livers of insulin resistant mice relative to control mice liver micrographs (Fig. S2, F).

To determine the impact of insulin resistance on the response to BpOmpW immunisation, groups of insulin resistant mice (Fig. S3, A) were immunized with one subcutaneous injection of adjuvant alone or with SAS-adjuvanted BpOmpW as before. After two weeks, the splenocytes of both groups were exposed to BpOmpW and immunophenotyped. Unexpectedly, CD45RB was not decreased in insulin resistance model (p = 0.6198; Fig. 4, A), despite the T cell activation markers, CD25 and CD44, being consistently elevated in both CD4^+^ and CD8^+^ T cells in BpOmpW immunised group with respect to the SAS control group (p < 0.0001, p < 0.0001, p = 0.0001, p = 0.0039, respectively; Fig. 4, B-E), which was comparable to non-insulin resistant mice. Insulin resistant BpOmpW immunised mice showed elevated levels of IL-2 in both CD4^+^ and CD8^+^ T cells following exposure to vaccine antigen relative to the splenocytes from adjuvant control mice (p = 0.0004 and p = 0.0003 respectively; Fig. 4, F-G) in contrast to the non-insulin resistant mice, which only showed upregulation of IL-2 in CD8^+^ T cells. Splenocyte cytokine responses from BpOmpW-immunised insulin resistant mice showed upregulated Th1 and Th17 responses as determined by the levels of IFN-γ and IL-17 in CD4 T cells, respectively (p < 0.0001; p < 0.0001; Fig. 4, H-I). In contrast to non-insulin resistant mice, Th2 response-associated IL-4 levels in CD4^+^ T cells were unaltered with respect to the control group (p = 0.2919; Fig. 4, J). Remarkably, after antigen restimulation, IFN-γ producing cytotoxic CD8 and NKT cells were also increased in BpOmpW immunised insulin resistant mice relative to the SAS control group (p = 0.0005, p = 0.0013; Fig. 4, K-L), with the exception of IFN-γ producing NK cells which remained unchanged (p = 0.6148; Fig. 4, M). Moreover, no significant changes were seen in IL-4 or IL-17-expressing NKT cells (Fig. 4, N-O). In contrast to non-insulin resistant mice, regulatory T cell levels remained unmodified (p value = 0.1729) in the splenocytes from BpOmpW immunised mice relative to SAS control splenocytes (Fig. 4, P). TNF in CD4 T cells was unchanged relative to the control group in the insulin resistance study (Fig. 4, Q), whereas TNF from CD8^+^ T cells was more abundant in BpOmpW group compared to control cells (p = 0.012; Fig. 4, R). We also analysed antibody responses and naïve and effector T cells in the insulin resistant model. We observed strong seroconversion (Fig. S3, B-D) and elevated effector T cells in the presence of BpOmpW (Fig. S3, E-I). Overall there were no remarkable changes in these immune parameters relative to the non-insulin resistant mice.

**Figure 4.**
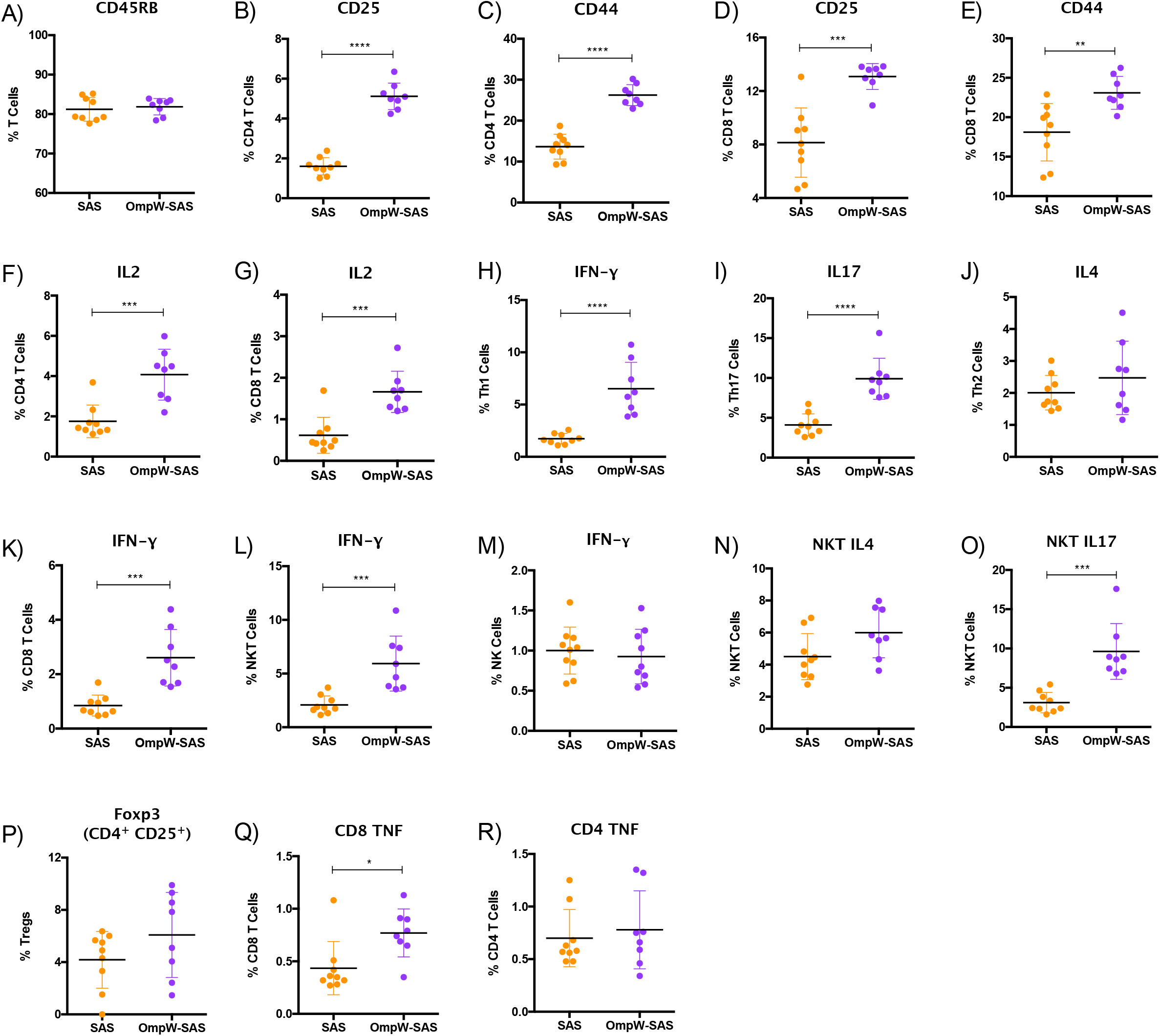
Insulin resistant mice mimicked with some changes the immune response to BpOmpW produced by non-insulin resistant mice. A) Percentages of CD45RB marker in total T cells. (B-E) Percentages of CD4 and CD8 T cells expressing CD25 and CD44 activation markers. F-G) Percentages of CD4 (F) and CD8 (G) T cells expressing IL-2 cytokine. H-J) Percentages of CD4 T cells expressing IFN-y (H), IL-17 (I) and IL-4(J). K-M) Percentages of CD8 (K), NKT (L) and NK (M) cells expressing IFN-y. N-O) Percentages of NKT cells expressing IL4 (N) and IL17 (O). P) Percentages of regulatory T cells (Tregs). Q-R) Percentages of CD4 (Q) and CD8 (R) T cells expressing TNF cytokine. SAS: splenocytes from adjuvant only immunised mice exposed to BpOmpW (Orange circles, Control). OmpW-SAS: splenocytes from SAS adjuvanted BpOmpW immunized mice re-exposed with BpOmpW in vitro (Purple circles, Treatment). Asterisks denote statistically significant differences according to two-tailed t-test. The significant levels are represented as follows: (p < 0.05, *); (p < 0.01, **); (p < 0.001, ***) (p < 0.0001, ****).

### BpOmpW elicited the IFN-γ recall responses in humanised HLA-DR and HLA-DQ transgenic mice

This and our previous work clearly demonstrate that BpOmpW antigen is immunogenic and elicits protective T cell responses in mice, thus as a precursor to its evaluation in human studies, we wanted to examine the responses in humanized HLA transgenic mice expressing different HLA alleles. The IFN-γ responses to the whole antigen were examined for HLA-DR1, HLA-DR4 and HLA-DQ8 alleles. In addition, a synthetic panel of overlapping peptides was generated covering the full coding sequence (Table 1) and these were also evaluated in humanised transgenic mice. Lymph nodes of HLA class II transgenic mice (HLA-DR1, HLA-DR4, and HLA-DQ8) that had been immunized with recombinant BpOmpW antigen showed strong recall responses to the whole BpOmpW antigen and to immunodominant T cell epitopes, P5, P7, P12, P21 as determined by IFN-γ ELISpot assay (Fig. 5). When probed further by priming of additional HLA-DR4 transgenic mice with P5 or P21, ELISpot analysis of drain lymph node cells confirmed that peptide 5 was a T-cell epitope. Subsequent priming of additional HLA-DQ8 mice with P7 confirmed that in HLA-DQ8 transgenics, P7 was a T-cell epitope (Fig. 5).

**Table 1.**
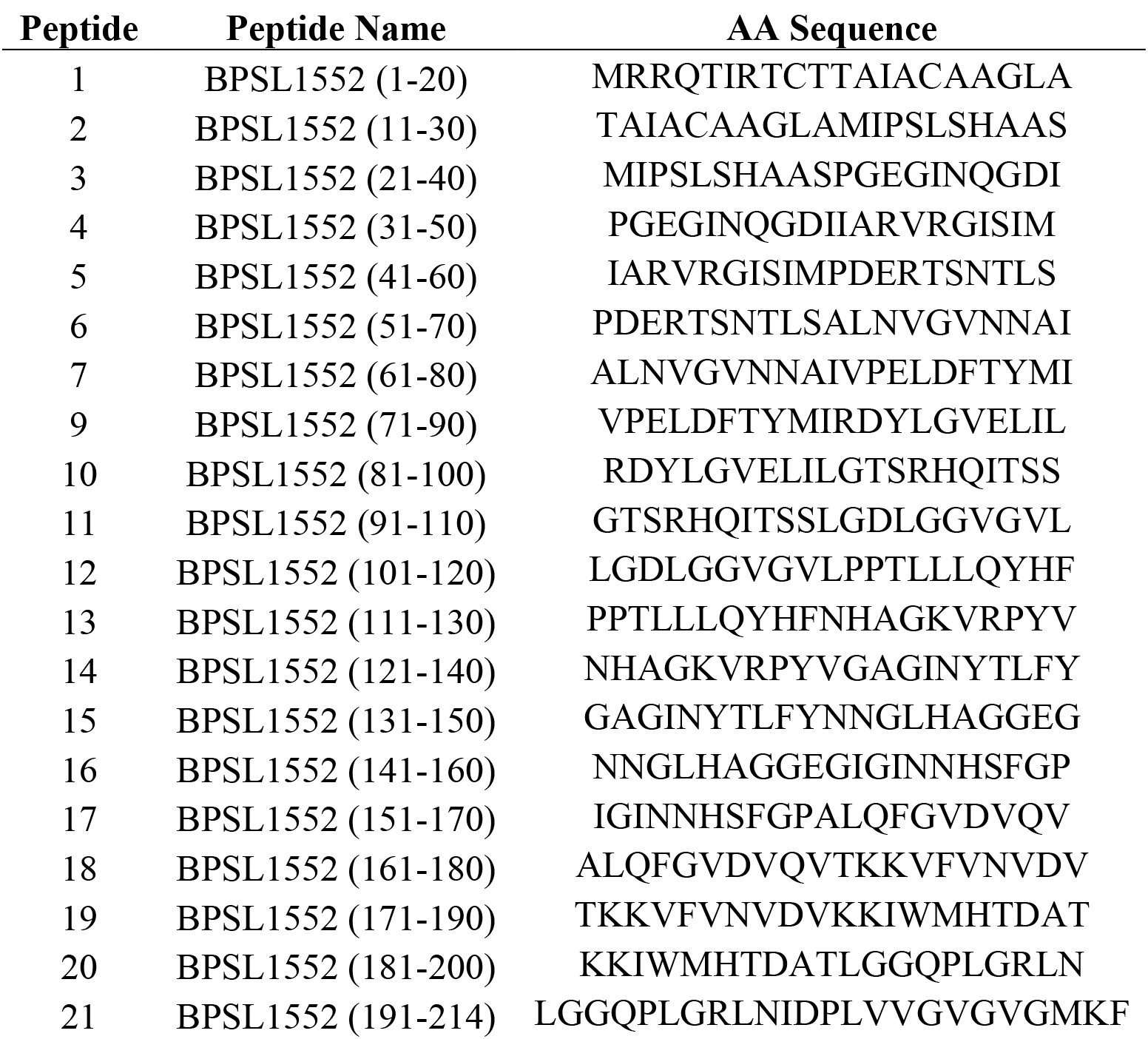
BpOmpW BPSL1552 peptide panel (accesion no. CAH35553.1)

**Figure 5.**
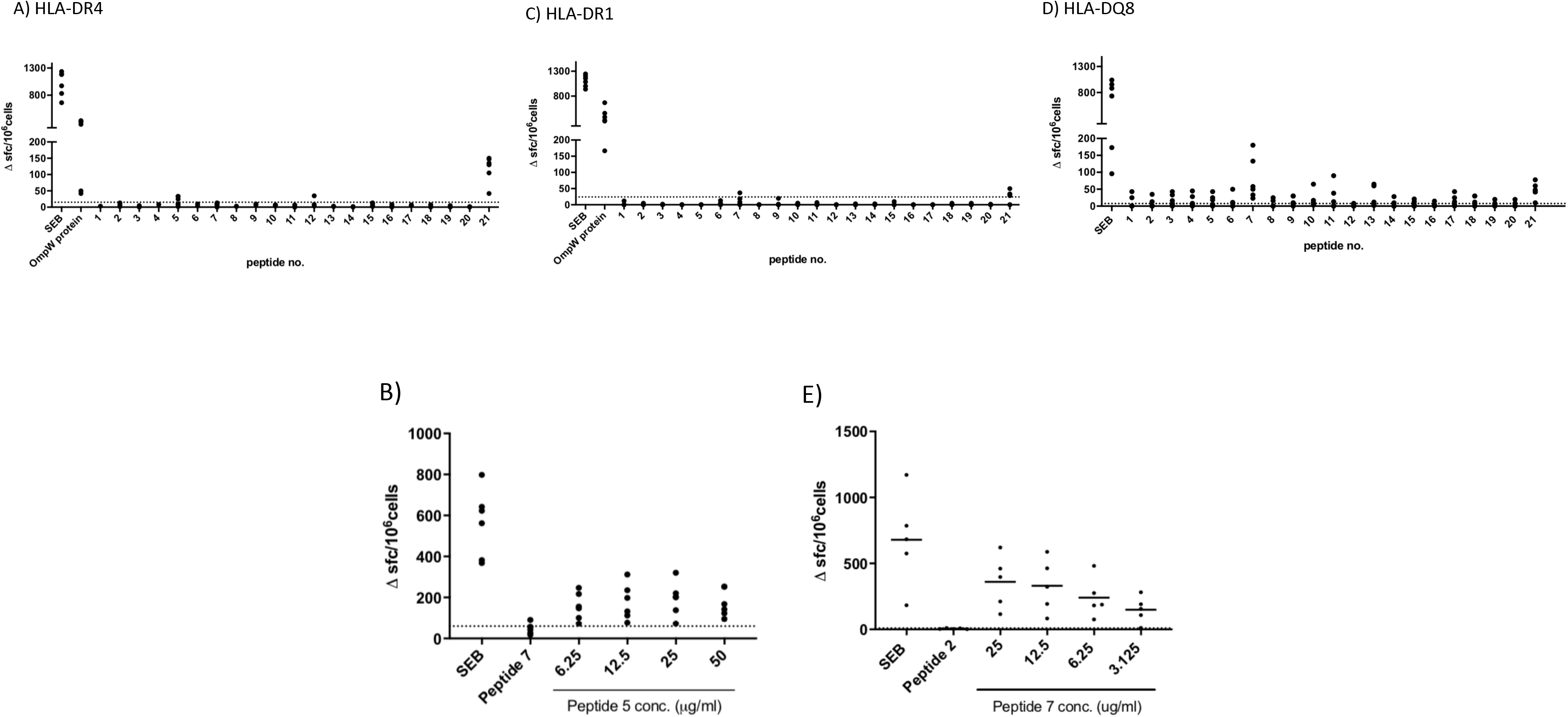
BpOmpW induced the production of IFN-γ in HLA-DR and HLA-DQ transgenic mice. Immunization of HLA-DR and -DQ transgenic mice highlights HLA class II determined immunodominant epitopes of BpOMpW. Mice transgenic for HLA-DR4, *n =*?, (A); HLA-DR1, *n =*? (B); and HLA-DQ8 (DQB1*0302), *n =*? (C) were primed with 25 μg rBpOmpW and draining lymph node cells were assayed with IFN-γ ELISpot in response to the indicated peptide at day 10. Data are plotted as SFCs per 10^6^ cells for individual mice. Responses to peptide were defined as positive if SFC > mean + 2 SD of the response in the absence of any antigen (shown as horizontal dotted line).

### BpOmpW induced T cell proliferation in human PMBCs

To examine the responses of human cells, we examined the T cell proliferation and IFN-γ responses of human PBMCs from donors in response to BpOmpW exposure. We interrogated human PBMCs from three different donors for CD3, CD4 and CD8 T cell markers following exposure to BpOmpW with and without adjuvant. T cell proliferation by BpOmpW was confirmed in all three different donor cell populations (Fig. 6 A, as proliferating cells for CD3, CD4 and CD8 populations were all elevated following exposure to the antigen with or without SAS adjuvant relative to the untreated control group (Fig. 6 A). T cells from donor 645 proliferated the most (Fig. 6 A) and cells from this donor were the only ones that showed significantly elevated IFN-γ responses in the presence of the antigen or when adjuvanted with respect to the control group (Fig. 6 B).

**Figure 6.**
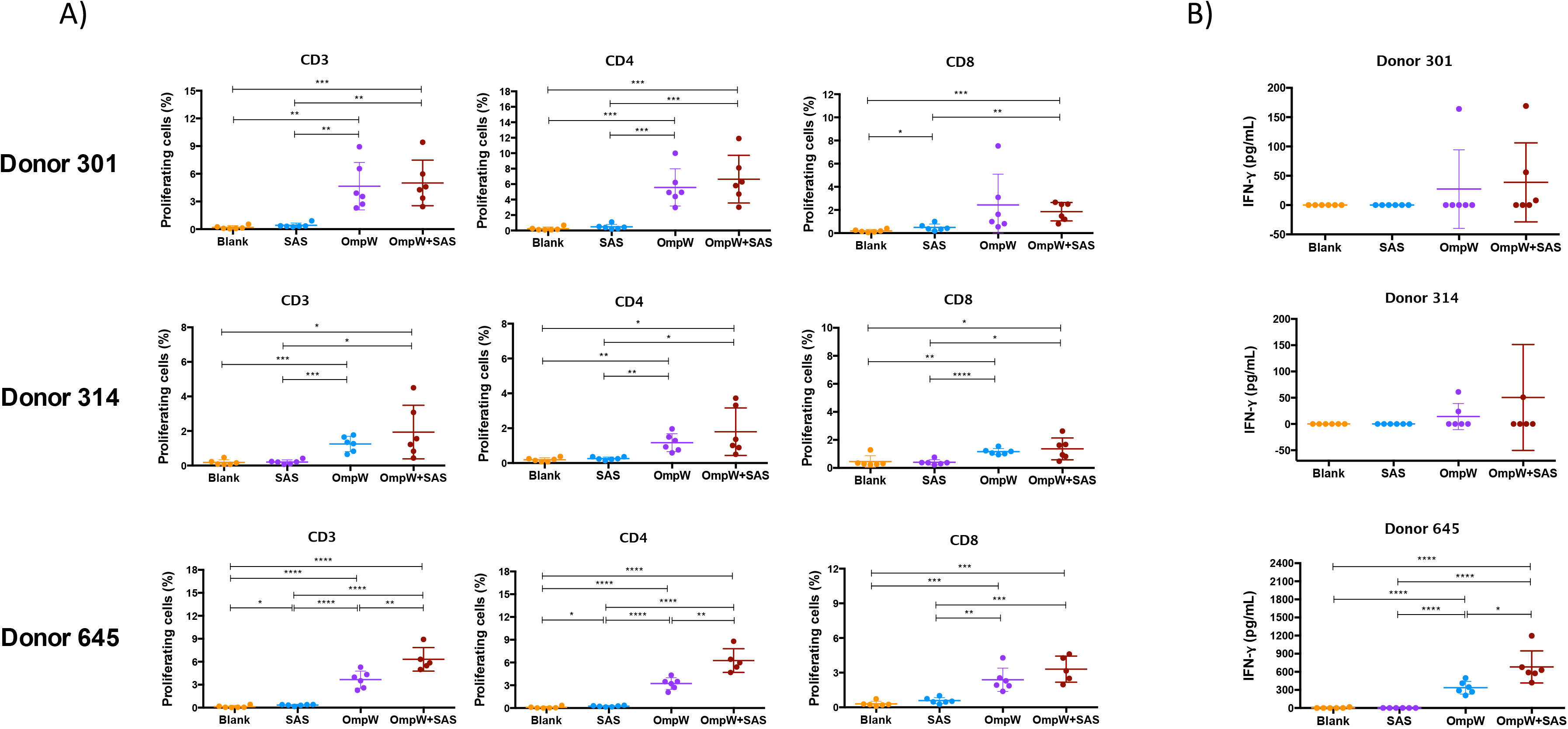
BpOmpW induced T cell proliferation in human PBMCs. A) Proliferation of CD3, CD4 and CD8 populations in three different donor PBMCs by BpOmpW or SAS-adjuvanted BpOmpW, measured as stimulation index. B) IFN-γ production (pg/mL) in the supernatant of the three proliferation assays. SAS: human PBMCs exposed to adjuvant alone (Orange circles, Control). OmpW: human PBMCs exposed to BpOmpW alone. OmpW-SAS: human PBMCs exposed to BpOmpW + SAS adjuvant. Asterisks mark significant differences according to two-tailed Student’s t-test. The significant levels are represented as follows: (p < 0.05, *); (p < 0.01, **); (p < 0.001, ***) (p < 0.0001, ****).

### BpOmpW is recognized by plasma from melioidosis survivors with Diabetes

We have shown that BpOmpW stimulates strong serological responses in mice, both in non-insulin resistant (Casey et al., 2016) and in insulin resistant mice. In order to examine whether the antigen can stimulate a humoral response in melioidosis patients with diabetes, we examined the presence of BpOmpW-specific antibodies in sera from people with diabetes that survived melioidosis infection (Fig. 7). We showed that the melioidosis survivor cohort with diabetes (M) had significantly higher BpOmpW-specific IgG responses than the healthy cohort with diabetes (p = 0.0289, D) and the non-endemic control (p = 0.0023, NE) groups, indicating that BpOmpW is specifically recognized and immunogenic in people with diabetes. No significant difference in IgG levels was seen between the melioidosis cohort and healthy household contacts (p = 0.0537, HH), most likely due to the endemic nature of melioidosis in Thailand.

**Figure 7.**
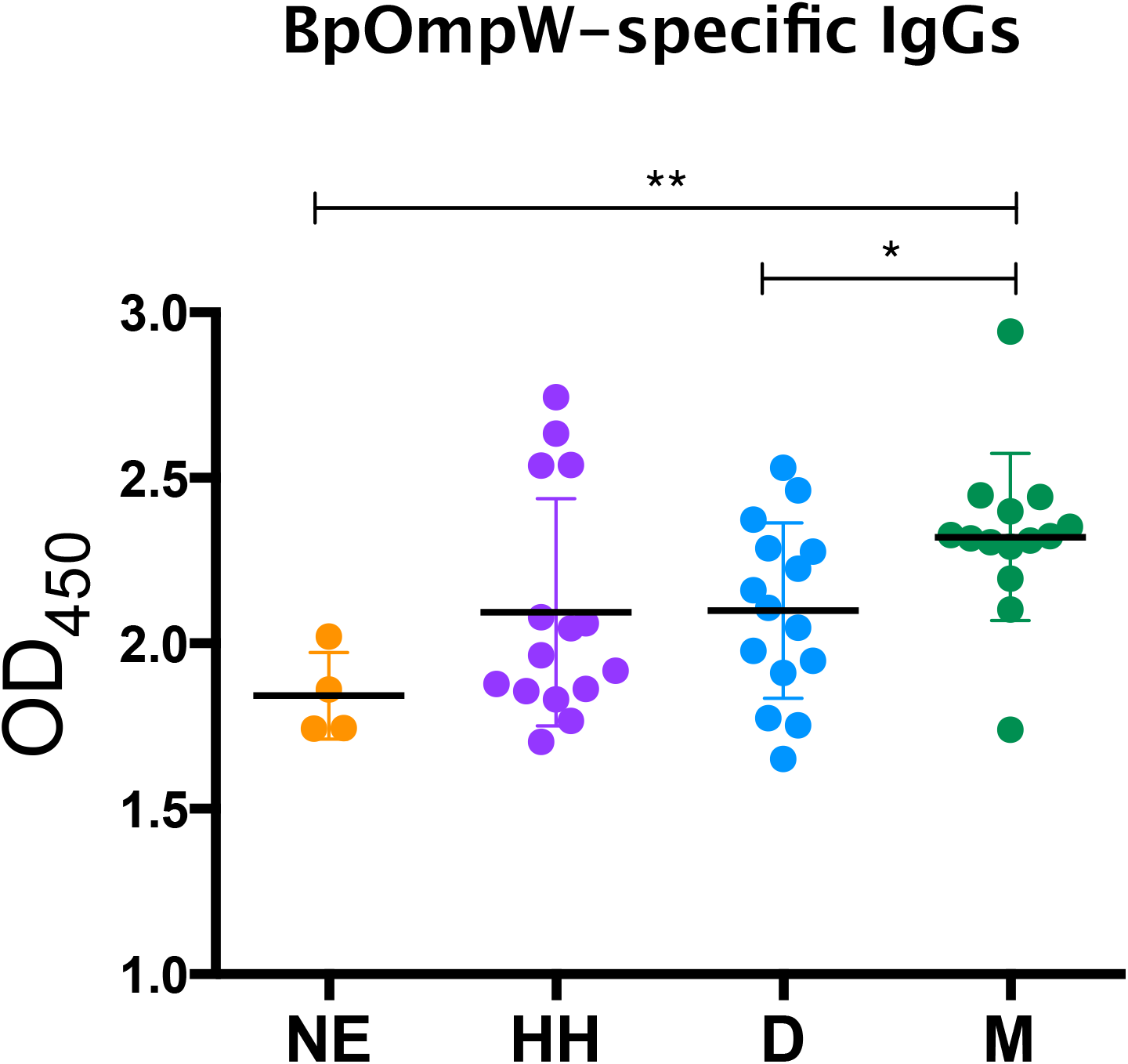
BpOmpW specific IgG responses in plasma from Melioidosis survivors. Detection by ELISA of BpOmpW-specific IgGs in different plasma from different cohorts (NE, Non-Endemic; HH: Healthy Householders; D, Healthy Diabetics; M: Melioidosis diabetics survivors). Asterisks mark significant differences according to two-tailed Student’s t-test. The significant levels are represented as follows: (p < 0.05, *); (p < 0.01, **); (p < 0.001, ***) (p < 0.0001,****).

## DISCUSSION

Melioidosis is a neglected tropical disease caused by the obligate intracellular bacterium *Burkholderia pseudomallei.* The continued emergence of this pathogen throughout tropical and subtropical regions (Wiersinga et al., 2012) together with the global increase of T2DM (Lin et al. 2020) is a cause of major concern in these areas and has accelerated the search of new remedies to combat this disease (Estes et al., 2010). Although the protective immune response against *B. pseudomallei* is not fully understood, it is well acknowledged that an effective vaccine should induce both T-cell and B-cell responses. Therefore, the elucidation of correlates of protection of the BpOmpW antigen in preclinical studies is essential in order to advance the development of an efficacious human T cell inducing vaccine against melioidosis.

The strong humoral responses elicited produced by BpOmpW in our earlier study (Casey et al. 2016) were also reproduced in this study in both non-insulin resistant and insulin resistant mice, confirming that the vaccine elicits potent specific antibody responses against BpOmpW. The splenocytes from BpOmpW immunized mice re-exposed to BpOmpW showed substantial activation and differentiation relative to control mice, as shown for example in the expression of CD25, and CD44 in both CD4 and CD8 T cells. T cell activation also leads to CD45RB regulation, an essential marker that determines the fate of T cells from naïve to memory cells. The generation of effector T cells that will undergo development to memory cells is an essential correlate of protection in determining the efficacy of vaccines (Robinson & Amara, 2005; Aravindhan et al., 2006). All effector subsets evaluated, i.e., effector CD4 and effectors CD8 were increased in BpOmpW immunized group, indicating that the antigen produces a robust effector recall response that is likely to lead to a memory response, a hallmark of an efficacious vaccine. The early immune responses to control melioidosis infection are predominantly mediated by IFN-γ in both CD4^+^ and CD8^+^ T cells, and as recently reported in NK and DN cells (Kronsteiner et al., 2019; Rongard et al., 2020). In this study, all these IFN-γ producing cell subsets were upregulated in the presence of the BpOmpW antigen, indicating that it elicits the required correlates of protection to combat the disease. Further, the vaccine elicited a mixed Th response, which is in line with the reported dominant Th2 and Th17 response at the initial stages of infection (Krishnananthasivam et al. 2017). The expression of the proinflammatory cytokine TNF is a typical murine host immune response to *B. pseudomallei* and was secreted in response to BpOmpW immunisation. At the same time, the elevated number of regulatory T cells following BpOmpW immunization observed in non-insulin resistant mice are required to reduce the excessive proinflammatory cascade of cytokines by bacterial lipopolysaccharides (Kessler et al., 2017). Finally, the fact that BpOmpW produced CD4hi and CD8hi populations suggest that, upon strong antigen stimulation, both populations of CD4hi and CD8hi may change from activated CD45RBlo CD44hi to naïve-like CD45RBhi CD44lo subsets in the BpOmpW group splenocytes as a regulatory mechanism against the strong antigen stimulation, although further analysis will be conducted to clarify this hypothesis. C57BL/6J mice are widely used as a model for chronic melioidosis (Hodgson et al., 2013 (Limmathurotsakul et al., 2015). HFD fed C57BL/6J mouse model has been previously used to study the immune responses impaired by DM in vaccine development (Haffer, 2012; Hodgson, 2013; Yoshida et al., 2020). In agreement with previous reports the mice gained weight and developed hyperglycemia and insulin resistance and showed the presence of lipid droplets in the liver within 12 weeks of HFD feeding. DM alters the adaptive immunity to infections (Hodgson et al., 2015), including melioidosis (Kronsteiner et al., 2019). The immune response to BpOmpW in insulin resistant mice was generally comparable to that seen in non-insulin resistant mice with some differences. BpOmpW re-exposure in insulin resistant mice maintained an activated T cell status and high IFN-γ recall responses from CD4, CD8, and NKT cells, which would be essential to protect people with diabetes from melioidosis. Non-insulin resistant mice showed a mixed Th1, Th2, Th17response, in response to BpOmpW immunisation in re-exposed splenocytes, in line with the Bcc-OmpW homolog examined previously (McClean et al., 2016). In contrast, insulin resistant mice showed Th1 and Th17 responses with no apparent Th2 response following BpOmpW immunisation. This is consistent with the fact that C57BL/6J strain preferentially differentiate to Th1 phenotype in response to HFD (Jovicic et al., 2015). Moreover, several studies also reported a delayed Th2 response in obese-allergic mice (Calixto et al., 2010; Silva et al., 2017; Esteves de Oliveira et al., 2019). Obesity, diabetes, and insulin resistance phenotypes produce proinflammatory cytokines that, in turn, are reported to downregulate regulatory T cells required to prevent the excessive inflammatory responses (Wagner et al., 2013; Cipolleta, 2014). In this work, regulatory T cells remained comparable to the control in the insulin resistance mice probably due to the stimulative effect of BpOmpW.

The T cell proliferation seen in different human PBMCs following BpOmpW exposure indicated that the vaccine antigen will also elicit T cell responses in humans. Moreover, the strong recall response of all HLA transgenic mice tested to the complete BpOmpW antigen indicates that the T-cell responses identified in humans are likely to translate to IFN-γ responses across diverse HLA isotypes. In contrast to other studies in which the identification of the candidate antigens was done on the basis of their reactivity against patient antisera, we looked for BpOmpW-specific antibodies in plasma from different cohorts, including survivors to melioidosis. The finding that BpOmpW also stimulated strong IgG responses in melioidosis survivors also indicates that it is likely to translate to a protective vaccine antigen to protect against this disease, although the ultimate test will require human trials. Overall the range of approaches used to elucidate whether the BpOmpW antigen elicits the necessary correlates of protection in humans strongly suggest that BpOmpW will elicit robust responses in humans and bring the vaccine closer to clinical trials.

## MATERIAL AND METHODS

### BpOmpW Expression and purification

The recombinant BpOmpW used in all experiments except ELISpot analysis of transgenic mice was expressed, purified and provided by Lionex GmBH in 20mM Ammonium bicarbonate. In the case of transgenic mouse studies, the pRSET_BpOmpW construct was transformed in BL21(DE3) cells and cultured in LB with 1M D-Sorbitol and 2.5mM glycine betaine for five days at 22°C. The His-tag fusion protein was then purified by nickel affinity chromatography with endotoxin-free PBS, 35mM imidazole, and 2% Triton X-100 and eluted in endotoxin-free PBS containing 250mM imidazole and 2% Triton X-100. The antigen was further purified by gel filtration chromatography. The affinity chromatography fraction containing the antigen (as identified by SDS-PAGE) was concentrated and loaded onto a HiLoad 16/600 GL Superdex 75 column (GE Healthcare) pre-equilibrated in endotoxin-free PBS using an AKTA chromatography system (GE Healthcare). Fractions with the protein of interest were pooled and the protein was concentrated and stored at – 80 °C until its use. Protein concentration was determined using the BCA protein assay kit (Thermo Fisher Scientific) and used for immunisation of transgenic mice and ELISpot assays.

### Ethics statement

All work involving animals was approved by University College Dublin Ethics Committee (AREC-19-13-McClean), and mice were maintained according to the regulations of the Health Products Regulatory Authority (Directive 2010/63/EU and Irish Statutory Instrument 543 of 2012) with the Authorisation number AE18982/P166.

For the human plasma samples from melioidosis patients and controls in Northeast Thailand, the study protocol was approved by the ethics committees of the Faculty of Tropical Medicine, Mahidol University (TMEC 12-014); Sunpasitthiprasong Hospital, Ubon Ratchathani (017/2559) and the Oxford Tropical Research Ethics Committee (OXTREC35-15). Blood samples were collected from in-patients with culture-confirmed melioidosis (M), diabetic patients (D), and healthy participants from the endemic areas who were household contacts of the melioidosis cases (HH) at Sunpasitthiprasong Hospital, Ubon Ratchathani, Thailand between 2015 and 2017.

All experiments used cryopreserved primary cells, i.e. PBMC, which were isolated from whole blood donated by healthy volunteers.

Whole blood was collected from healthy donors as described in the ethical protocol IXP-003_V1 (Belgian registration number B707201627607) or protocol IXP-004_V1 (The Netherlands; Reg. Nr. NL57912.075.16). All blood samples were tested and found negative for HBV, HCV and HIV. PBMC were separated from the blood by density gradient centrifugation and subsequently cryopreserved in FBS, supplemented with 10% dimethyl sulfoxide, by controlled rate freezing. The PBMCs were kept in cryogenic storage (−180°C) until use.

### Immunisation of C57BL/6J mice for immunophenotyping

Male C57BL/6J mice were used in these studies and were given free access to food and water and subjected to a 12h light/dark cycle. Groups of C57BL/6J male mice were immunized subcutaneously with 50μg BpOmpW in Sigma Adjuvant System (SAS, Sigma) or SAS adjuvant alone as negative control. Two weeks later, mice were humanely killed by sedation with isoflurane followed by CO_2_ exposure and, blood removed by cardiac puncture for serological analysis. Spleens were also processed for splenocyte restimulation with the vaccine antigen.

### Determination of BpOmpW-specific IgG isotypes by ELISA

Microtitre plates were coated with purified BpOmpW in sodium bicarbonate buffer (pH 9.4) at 4°C overnight. Coating solution was removed, and plates blocked with 10% FBS solution in PBS at room temperature for 1h. Wells were washed three times with 0.05% Tween 20 in PBS using a plate washer. Serum samples were serially diluted (5-fold) in PBS containing 10% FBS and 100μl added to wells in triplicates at RT for 2h. Plates were washed three times as described above with PBS 0.05% Tween 20 before the addition of anti-mouse IgG, IgG1 or IgG2a-HRP conjugated antibodies (ab97023, ab97240, ab97245, respectively from Abcam) at RT for one hour. Then, TMB substrate was added and incubated until the colour developed. Reactions were stopped with 2M sulfuric acid and plates read at 450nm.

For the detection of BpOmpW-specific IgG in human plasma the same protocol was applied using anti-human IgG antibody (ab6858, abcam) in place of anti-mouse IgG antibodies.

### Splenocyte Restimulation with BpOmpW antigen

Splenocytes were extracted from the spleens using Ammonium-Chloride-Potassium (ACK) lysing buffer to remove red blood cells. Cells from BpOmpW immunized or SAS adjuvant only treated control mice were then counted automatically (Invitrogen). One million cells were plated per well, and stimulated for 60h with 50μg/mL BpOmpW in 96-well-plate using 10% FBS RPMI medium and P/S. Between five and six hours before harvesting the cells, 5μg/mL Brefeldin A was added to block cell trafficking in order to increase the accumulation of intracellular cytokines. Cells were collected by centrifugation and stained for flow cytometry.

### Flow Cytometry

The cells were incubated with Fc Block (anti-CD16/CD32; BD BioSciences) for 5 minutes (TEMP) and labeled with Viakrome 808 (Beckman Coulter) and fluorochrome-labeled antibodies against CD4, CD49b, CD45RB, CD8a, CD25, CD44 and CD3 surface markers (BD Biosciences) for 30 minutes (TEMP). Then, intracellular IL-2, IL-4, IFN-γ, IL-9, TNF, and Foxp3 (BD Bioscience) were also analyzed with a BD Cytofix/CytoPerm^™^/Fixation/Permeabilization Solution Kit (BD Biosciences) according to the manufacturer instructions. The gating strategy and the corresponding FMOs for each gate are shown in Figure S3-S9.

### Polygenic Insulin Resistant Mouse Model

Seventy C57BL/6J male mice were fed with D12492i rodent diet comprised of 60% kcal from fat (Research Diets, Inc) or regular chow starting at 6-8 weeks of age until they were humanely killed. Insulin resistance was determined at 8 and 12 weeks by fasting the mice for 6h at which time 0.5 units/Kg of insulin was administered subcutaneously. In order to alleviate any pain, EMLA cream was applied to the whole tail 10 minutes before blood was sampled from the tail vein using a 27-gauge needle, a drop of blood extracted at 15, 30, 45 & 60 minutes, and glucose levels measured by AlphaTRAK^®^ 2 Blood Glucose Monitor. To characterise the model, we collected the spleen, liver, pancreas, kidney from the mice and H&E stain was applied to tissue cut sections. The time required for individual mice to develop insulin resistance varied and only mice that showed insulin resistance were considered eligible to be randomly selected for immunization studies.

### ELISpot analysis of IFN-γ recall response to antigen or peptides

This study used HLA class II transgenic mouse lines for the alleles HLA-DR1 (DRB1*0101), HLA-DR4 (DRB1*0401) and HLA-DQ8 (DQB1*0302), which were in each case maintained in the context of a homozygous knockout for murine H2-Ab. Mice were maintained in individually ventilated cages and were used in experiments as age- and sex-matched, young adults. For CD4 T cell epitope mapping studies, mice were primed in one hind footpad with 25mg antigen emulsified in Hunters Titermax Gold adjuvant (Sigma-Aldrich). At day 10, the draining popliteal lymph node was removed and disaggregated into a single-cell suspension for ELISpot assays. The frequency of cells producing IFN-γ in responses to antigen was quantified with ELISpot (Diaclone; 2B Scientific, Oxon, U.K.) performed in HL-1 serum-free medium (BioWhittaker; Lonza, Slough, U.K.), supplemented with L-glutamine and penicillin–streptomycin (Life Technologies, Paisely,U.K.). Cells (23105) plus antigen were added to wells and plates and were incubated for 72 h at 37°C with 5% CO_2_. Unless otherwise indicated, peptide was added to wells at a final concentration of 25mg/ml. Spots were counted on an automated ELISpot reader (Autoimmun Diagnostika, Strasbourg, France). Response frequencies were expressed as **Δ**SFC/10^6^cells, with the presence of an epitope being confirmed when the majority of mice in the immunized group responded with a magnitude greater than the mean number of spot-forming cells (SFCs) in medium only control + 2 SD. Mean + 2 SD background SFC for murine ELISpot data are indicated in each case by a dotted line on the figures. The ELISpot background ranges (per 10^6^ cells) were 0 to 30 SFC.

### Statistical Analysis

Results are presented as means ± SE unless otherwise stated. The differences in means between groups were tested using a T-test using GraphPad Prism, version 7. A p-value < 0.05 was considered statistically significant.

## ACKNOWLEDGEMENTS

Supported by a Wellcome Trust Innovator Award to SMcC (209274/Z/17/Z). JTC is recipient of the Basque Government Postdoctoral Fellowship. CQ is a recipient of funding from UCD^2^ Transatlantic One Health Alliance.

## AUTHOR CONTRIBUTION

Performed the experiments: JTC, LB, CQ, CR, DB, NC, MOM, EM Analyzed the data: JTC, JA, AB, DA, RB, SMcC. Designed and supervised the experiments: JTC, JA, AB, DA, RB, JA, SG, SMcC. Wrote the manuscript: JTC, SMcC.

## ETHICS DECLARATION

