## Supplementary figures and images for "BpOmpW Antigen Stimulates the Necessary Immune Correlates of Protection Against Melioidosis"

### Supplemental figures

#### Figure S1

**Non insulin resistant mice:**

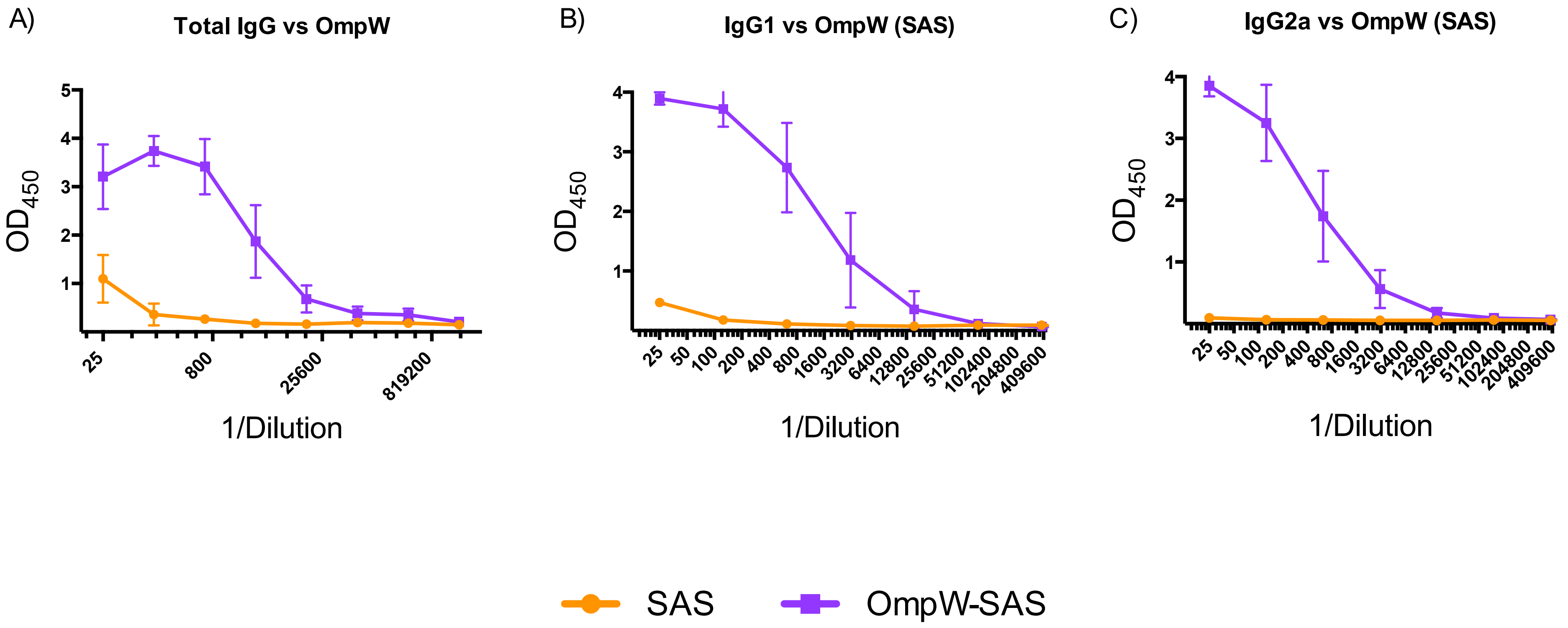

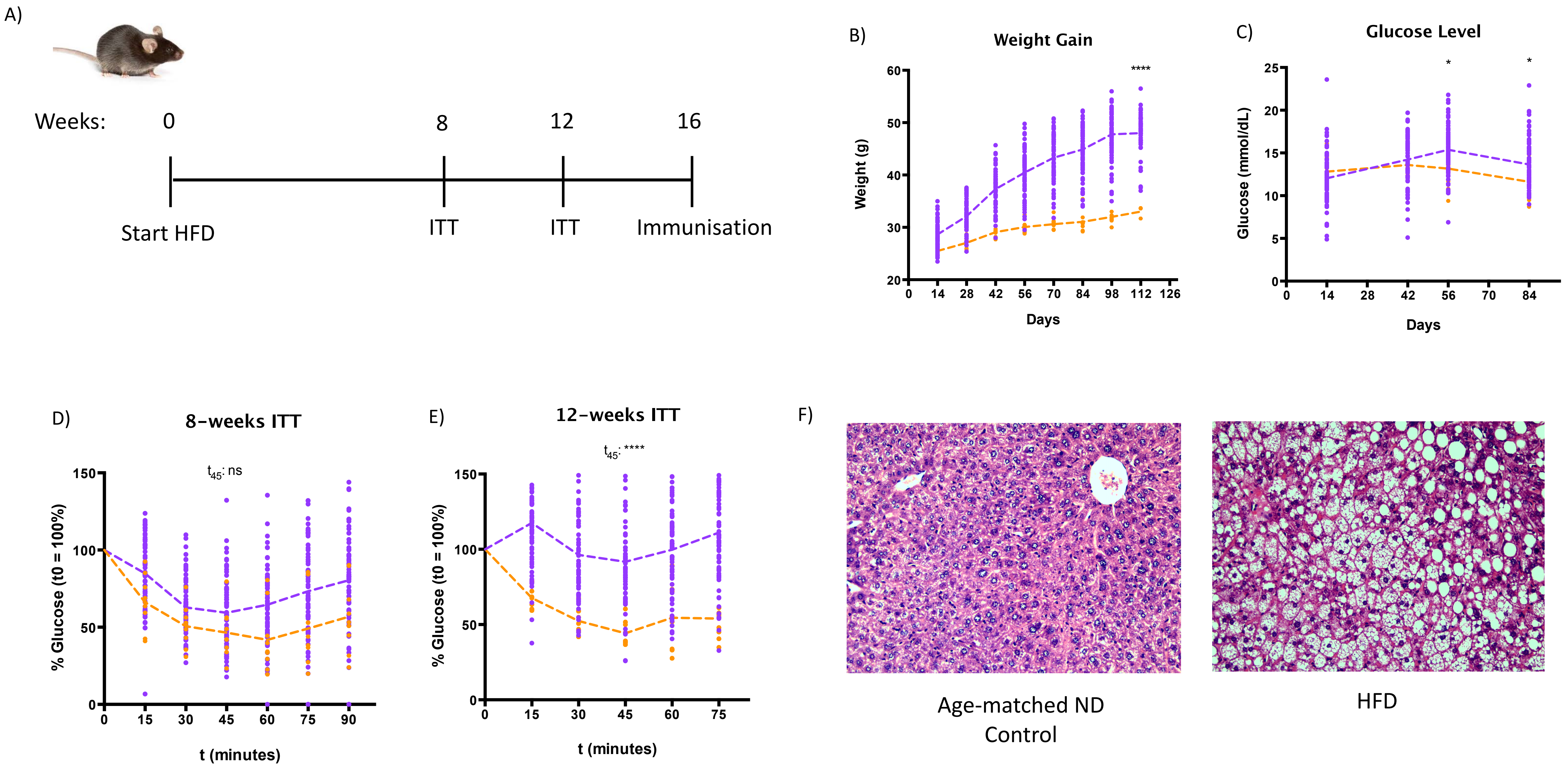

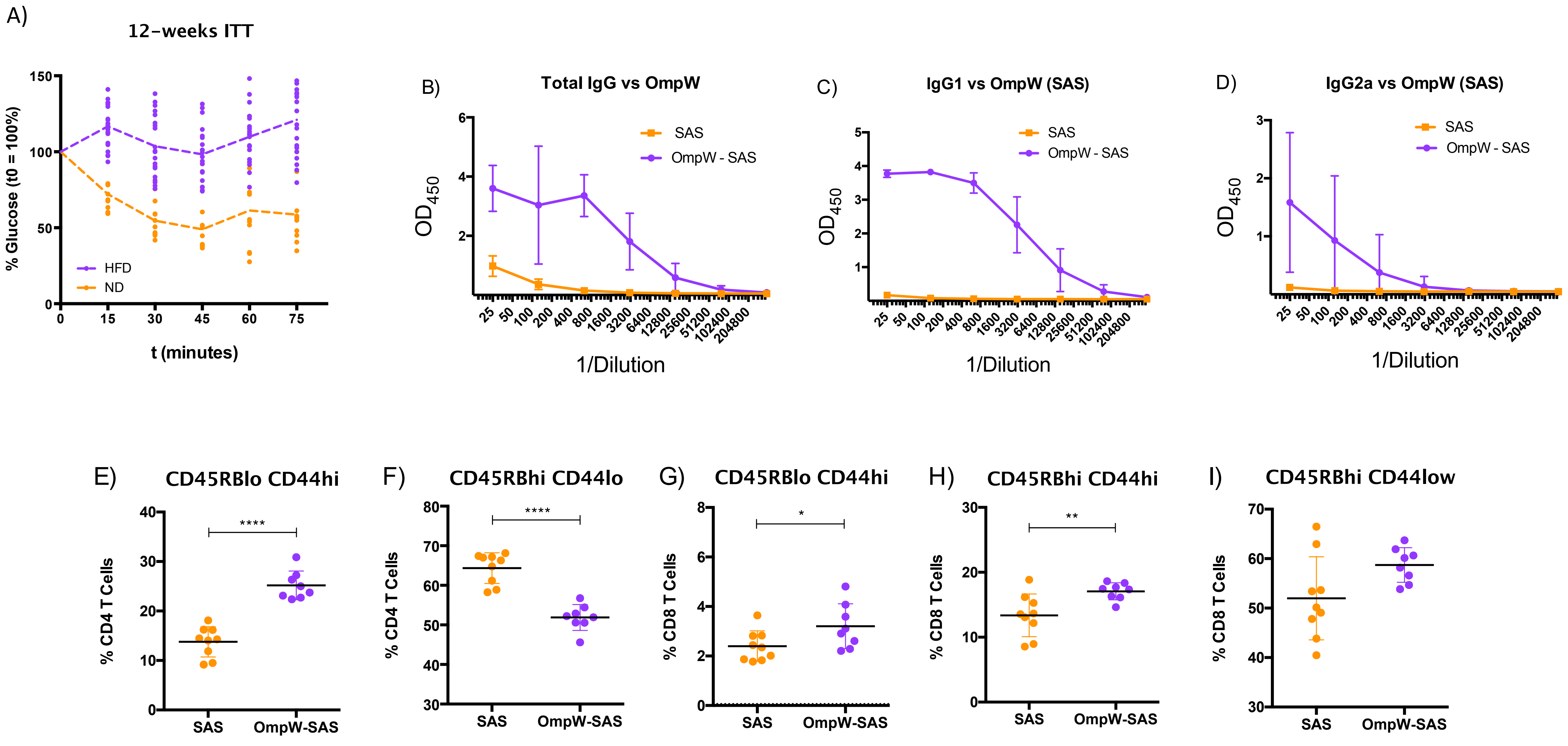

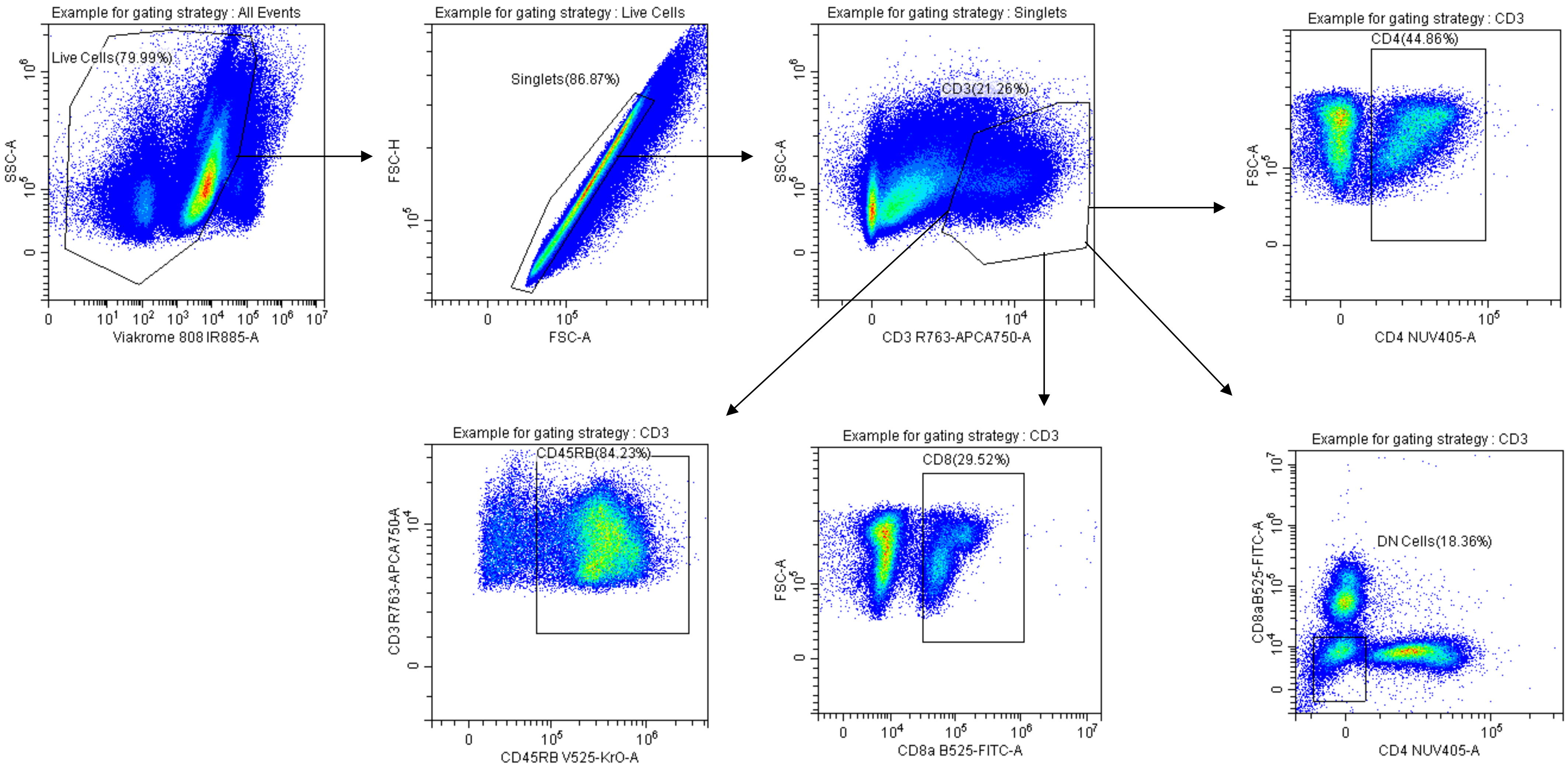

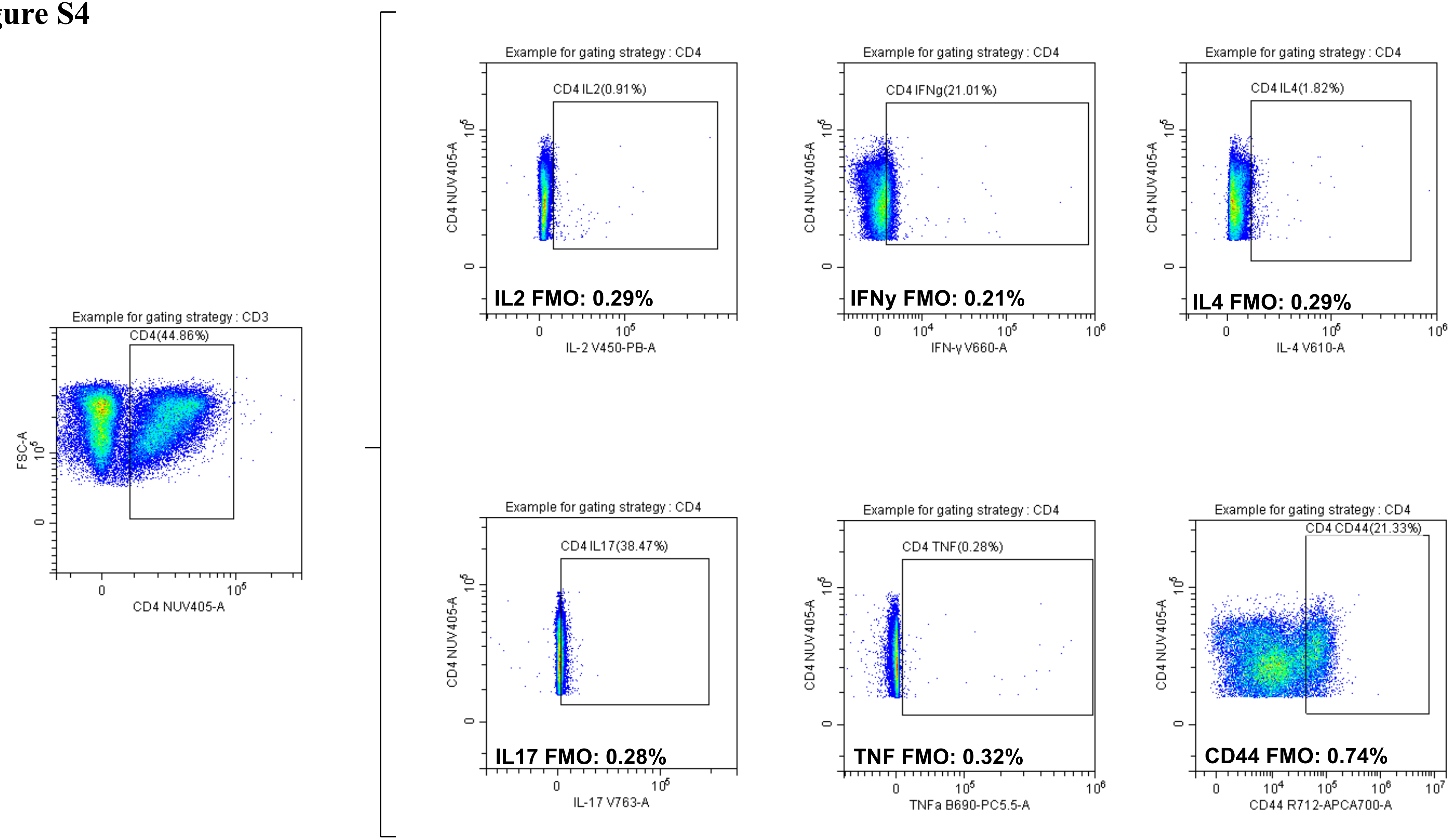

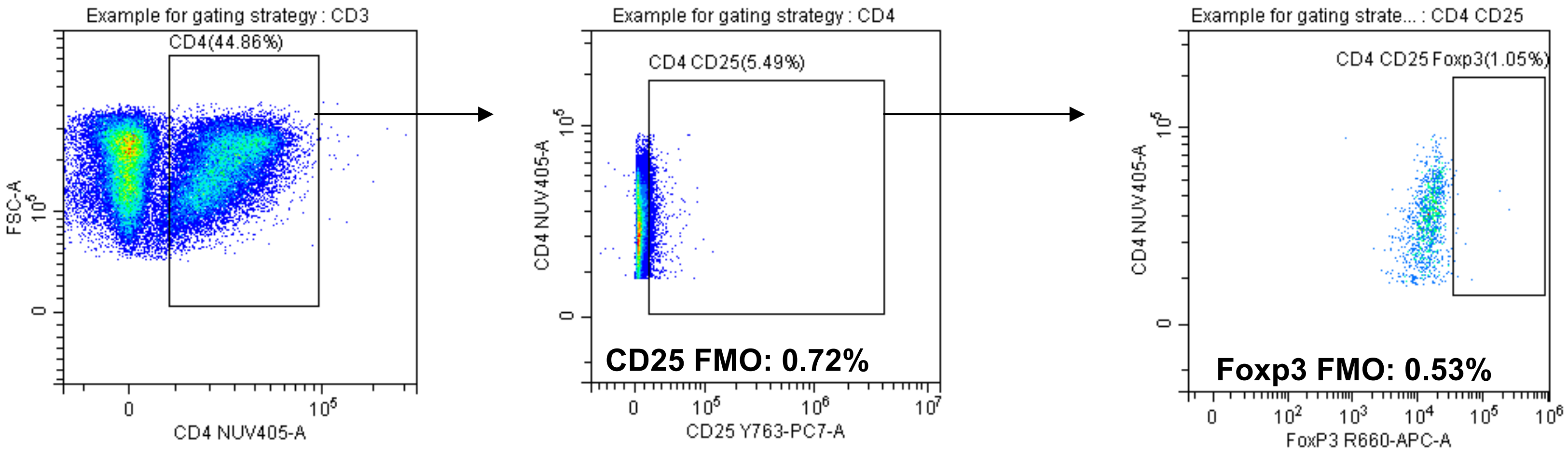

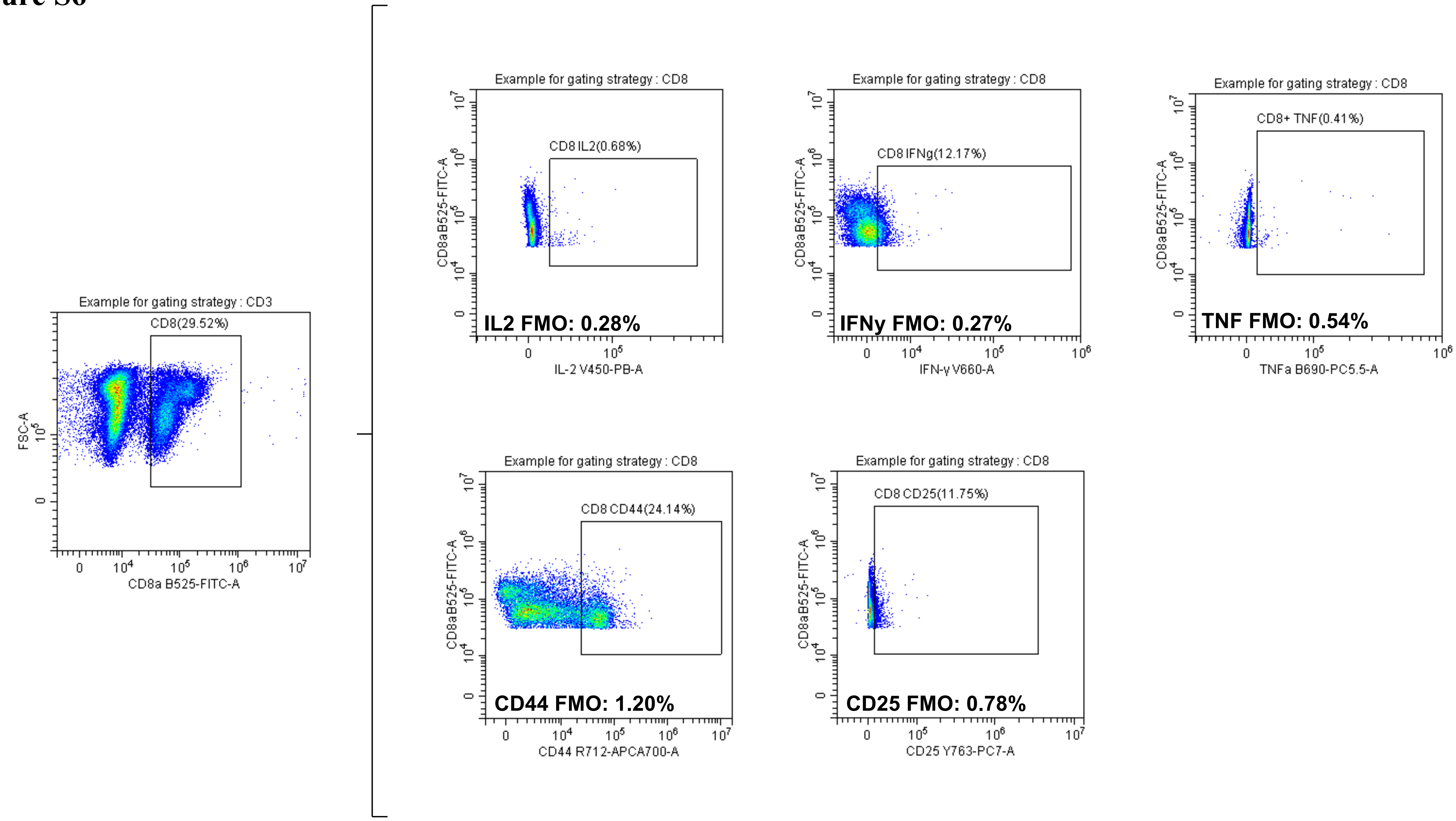

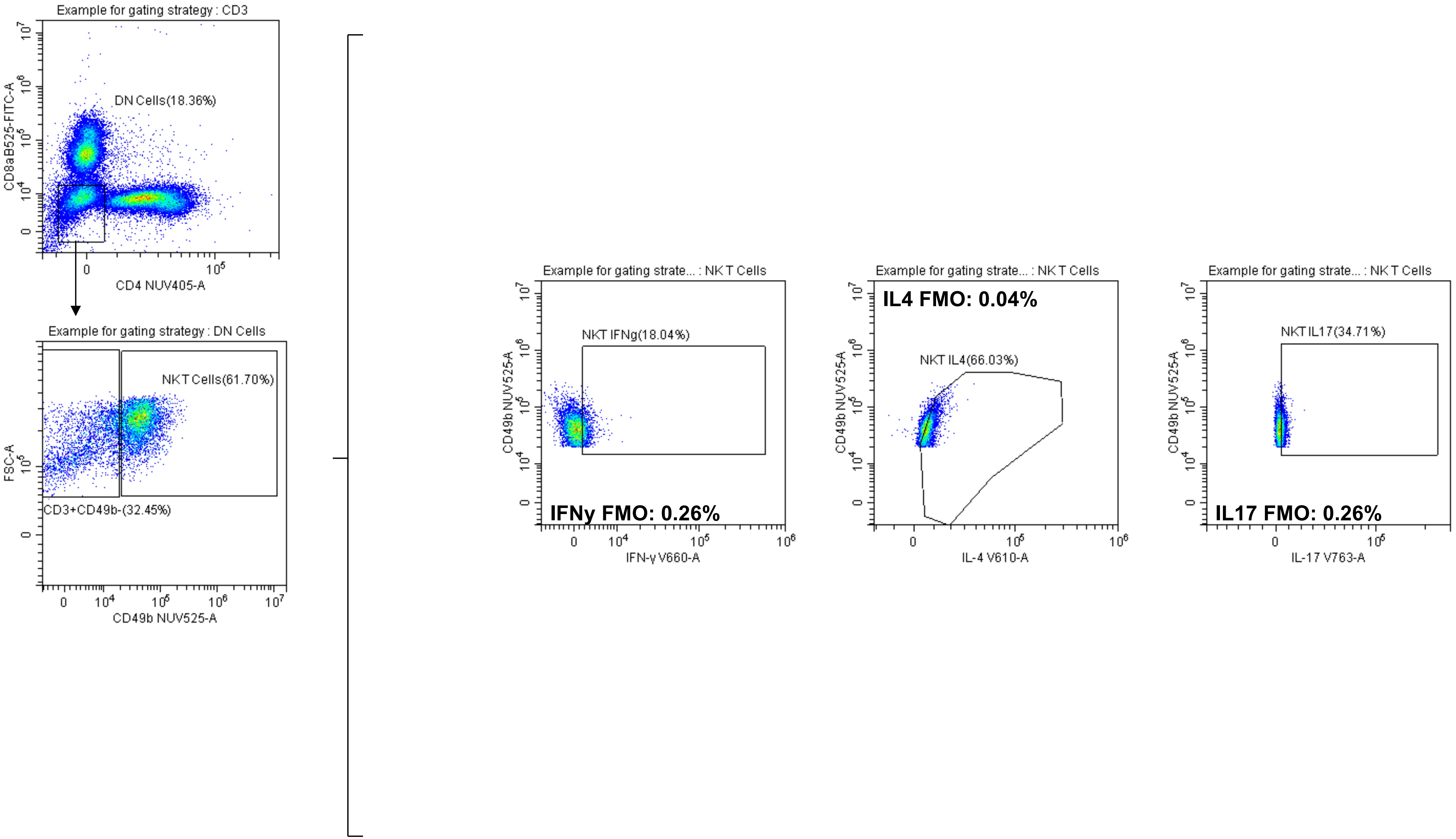

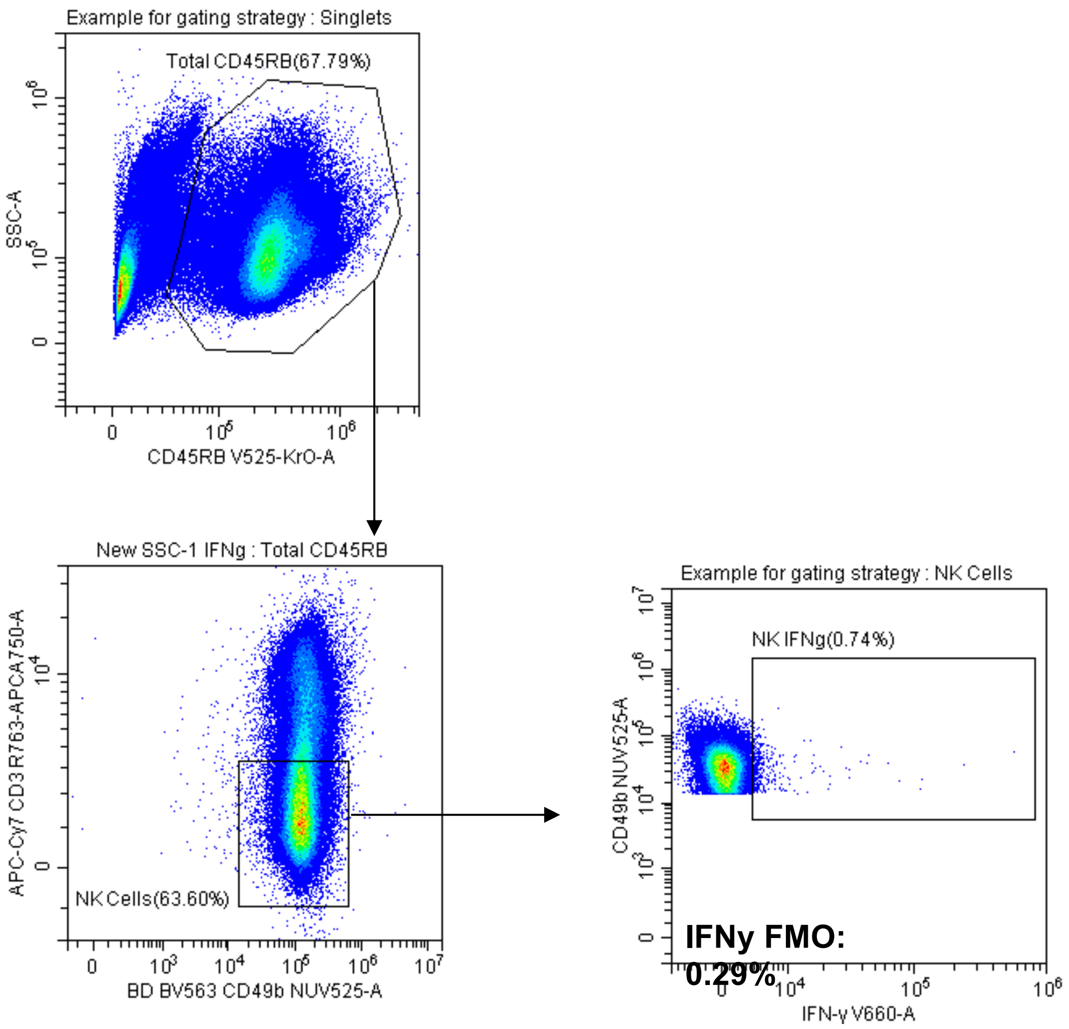

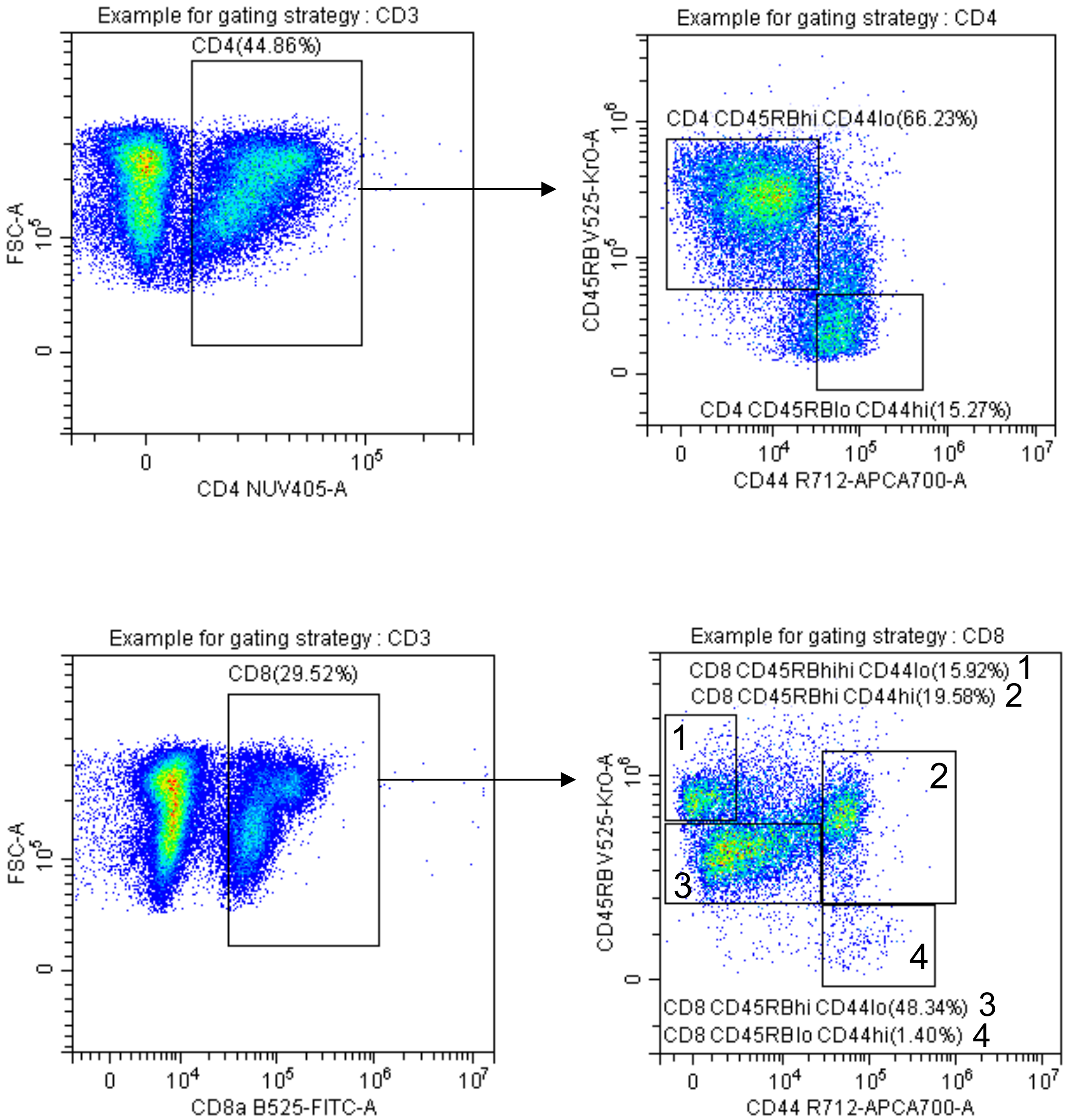
